# *Sellimonas caecigallum* sp. nov., description and genome sequence of a new member of the Sellimonas genus isolated from the cecum of feral chicken

**DOI:** 10.1101/714204

**Authors:** Supapit Wongkuna, Sudeep Ghimire, Linto Antony, Surang Chankhamhaengdecha, Tavan Janvilisri, Joy Scaria

**Affiliations:** Department of Veterinary and Biomedical Sciences, South Dakota State University, Brookings, SD, USA; South Dakota Center for Biologics Research and Commercialization, SD, USA; Department of Biochemistry, Faculty of Science, Mahidol University, Bangkok, Thailand; Department of Biology, Faculty of Science, Mahidol University, Bangkok, Thailand

## Abstract

An obligately anaerobic, non-motile, Gram-positive coccobacillus strain SW451 was isolated from pooled cecum contents of feral chickens. Comparative analysis based on 16s rRNA sequence showed that strain SW451 had 95.24% nucleotide sequence similarity to *Sellimonas intestinalis* BR31^T^, the closest species with a valid taxonomy. The genome of SW451 is 2.67 Mbp with 45.23 mol% of G+C content. The major cellular fatty acids were C_16: 0_, C_14: 0_ and C_16: 0_ DMA. Based on taxonogenomic, physiological, and biochemical analysis, the strain SW451 represents a new species of the genus *Sellimonas*, for which the name *Sellimonas caecigallum* sp. nov. is proposed. The type strain of *Sellimonas caecigallum* is SW451 (=DSM 109473^T^)

## Introduction

The chicken gut contains a diverse microbial community which is dominated by the phyla Firmicutes, Actinobacteria, Proteobacteria and Bacteroidetes (1–3). Gut commensal bacteria play an important role in host metabolism, immune system and pathogen protection (4, 5). Recently, gut microbiota has been studied using culture-independent and culture-dependent techniques. Sequencing-based analysis of the gut microbiome is useful for identifying dynamic changes in microbiota composition, while culturing is used to access biological functions of individual microbiota members and their interactions (6–8). However, the cultivation and isolation of several bacterial species from gut microbiota remains a daunting problem. Here, we report the cultivation and characterization of a new bacterial species from the cecum of feral chicken. The proposed *Sellimonas caecigallum* sp. nov. strain SW451 (=DSM 109473) is a new member of genus Sellimonas.

## Isolation, growth conditions and strain identification

Strain SW451 was isolated from cecum of feral chicken by culturing in strict anaerobic conditions in a Coy Lab anaerobic chamber containing 85% nitrogen, 10% hydrogen and 5◻% carbon dioxide. Modified Brain Heart Infusion (BHI-M) medium, which contained 37 g/L of BHI, 5 g/L of yeast extract, 1 ml of 1 mg/mL menadione, 0.3 g of L-cysteine, 1 mL of 0.25 mg/L of resazurin, 1 mL of 0.5 mg/mL hemin, 10 mL of vitamin and mineral mixture, 1.7 mL of 30 mM acetic acid, 2 mL of 8 mM propionic acid, 2 mL of 4 mM butyric acid, 100 μl of 1 mM isovaleric acid, and 1% of pectin and inulin was used for strain culturing and maintenance. Genomic DNA of the strain SW451 was extracted using DNeasy Blood & Tissue kit (Qiagen), according to the manufacturer’s instructions. 16S rRNA gene sequences was amplified using universal primer set 27F (5’-AGAGTTTGATCMTGGCTCAG-3’) and 1492R (5’-ACCTTGTTACGACTT- 3’) (9, 10). The PCR amplicon was sequenced using Sanger dideoxy method (ABI 3730XL; Applied Biosystems). The 16S rRNA gene sequence of SW451 was compared to closely related strains obtained from the GenBank (www.ncbi.nlm.nih.gov/genbank/) and EZtaxon databases (www.ezbiocloud.net/eztaxon) (11). Phylogenetic analysis was conducted using MEGA7 software (12). Multiple alignments were generated using the CLUSTAL-W algorithm (13). Reconstruction of phylogenetic trees was carried out using the maximum-likelihood (ML) (14), maximum-parsimony (MP) (15), and neighbour-joining (NJ) (16) methods. The distance matrices were generated using Kimura’s two-parameter model. Bootstrap resampling analysis of 1000 replicates was performed to estimate the confidence of tree topologies. Based on the results of 16S rRNA gene sequencing, the closest taxa of SW451 were *Sellimonas intestinalis* DSM 103502^T^ (95.24◻% similarity) and *Clostridium nexile* KCTC 5578^T^ (94.47%), followed by *Merdimonas faecis* KCTC 15482^T^ (94.34%) and *Faecalimonas umbilicata* DSM 103426^T^ (94.08%). The results of phylogenetic analysis showed that SW451 was clustered with the *Sellimonas* clade and it represents a new member of the genus *Sellimonas* (Fig. 1).

**Fig. 1.**
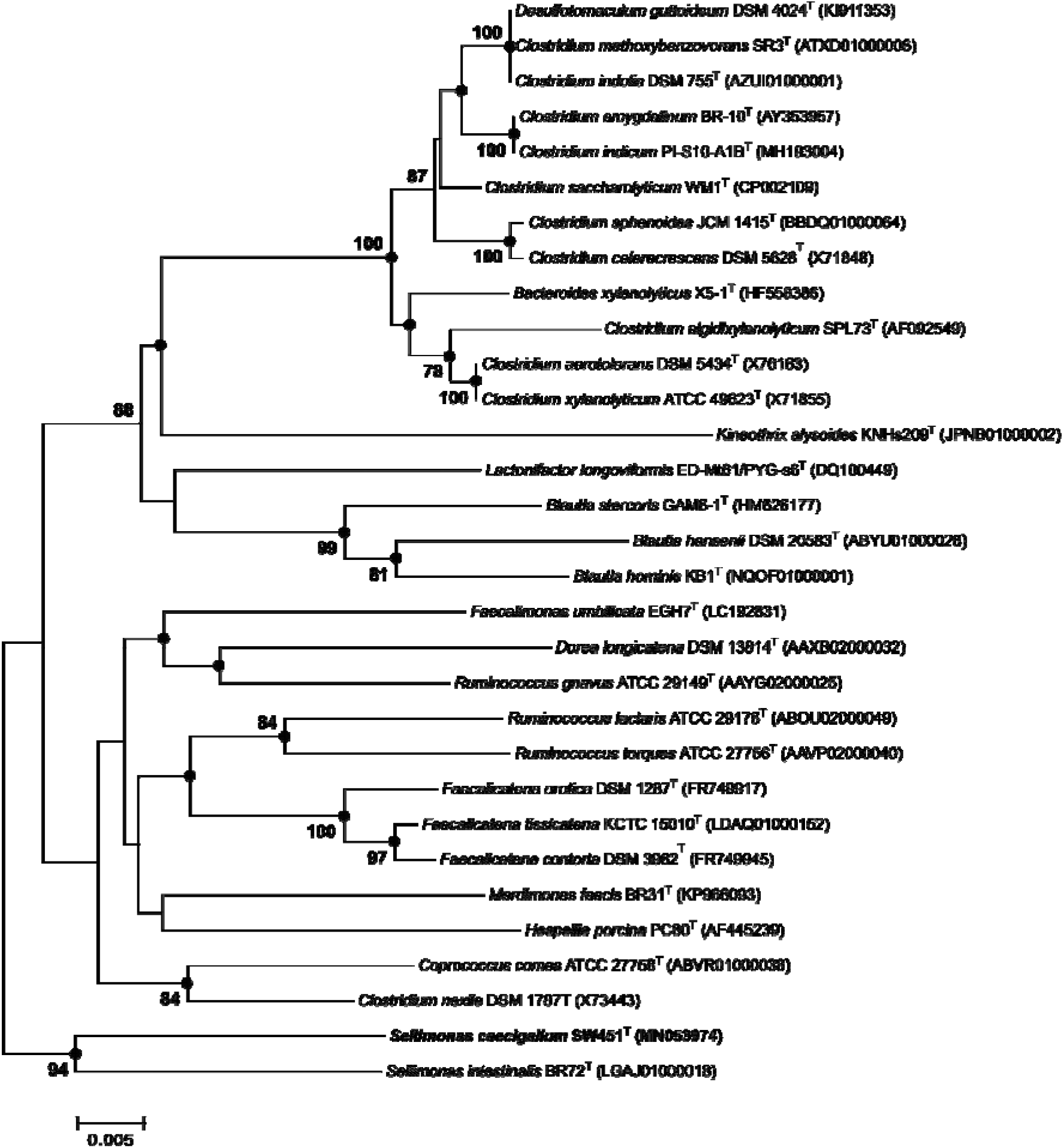
Neighbor-joining tree based on 16S rRNA gene sequences, showing the phylogenetic position of Sellimonas caecigallum within closely related taxa in the genus Sellimonas of Clostridium cluster XIVa. GenBank accession numbers of the 16S rRNA gene sequences are given in parentheses. Black circles indicate that the corresponding branches were also recovered both by maximum-likelihood and maximum parsimony methods. Number at nodes are shown as percentages of bootstrap greater than 70 %. Bar, 0.005 substitutions per nucleotide position.

## Genome properties and comparison

The whole genome sequencing of strain SW451 was performed using Illumina MiSeq sequencer with Illumina V3 2 x 250 chemistry. The reads were assembled using Unicycler that builds an initial assembly graph from short reads using the *de novo* assembler SPAdes 3.11.1 (17). The quality assessment for the assembly was performed using QUAST (18).The draft genome of strain SW451 has total length of 2.67 Mbp. The genomic G+C content of SW451 was 45.13◻mol%. Genome features of the strain was distinct from other related strains (Table 3). In addition, the average nucleotide identity (ANI) was calculated between SW451 and the closest related strains using OrthoANI software (19). The ANI values were significantly less than the proposed ANI cutoff of 95– 96◻% (20), demonstrating strain SW451 as a novel species (Fig. 2).

**Fig. 2.**
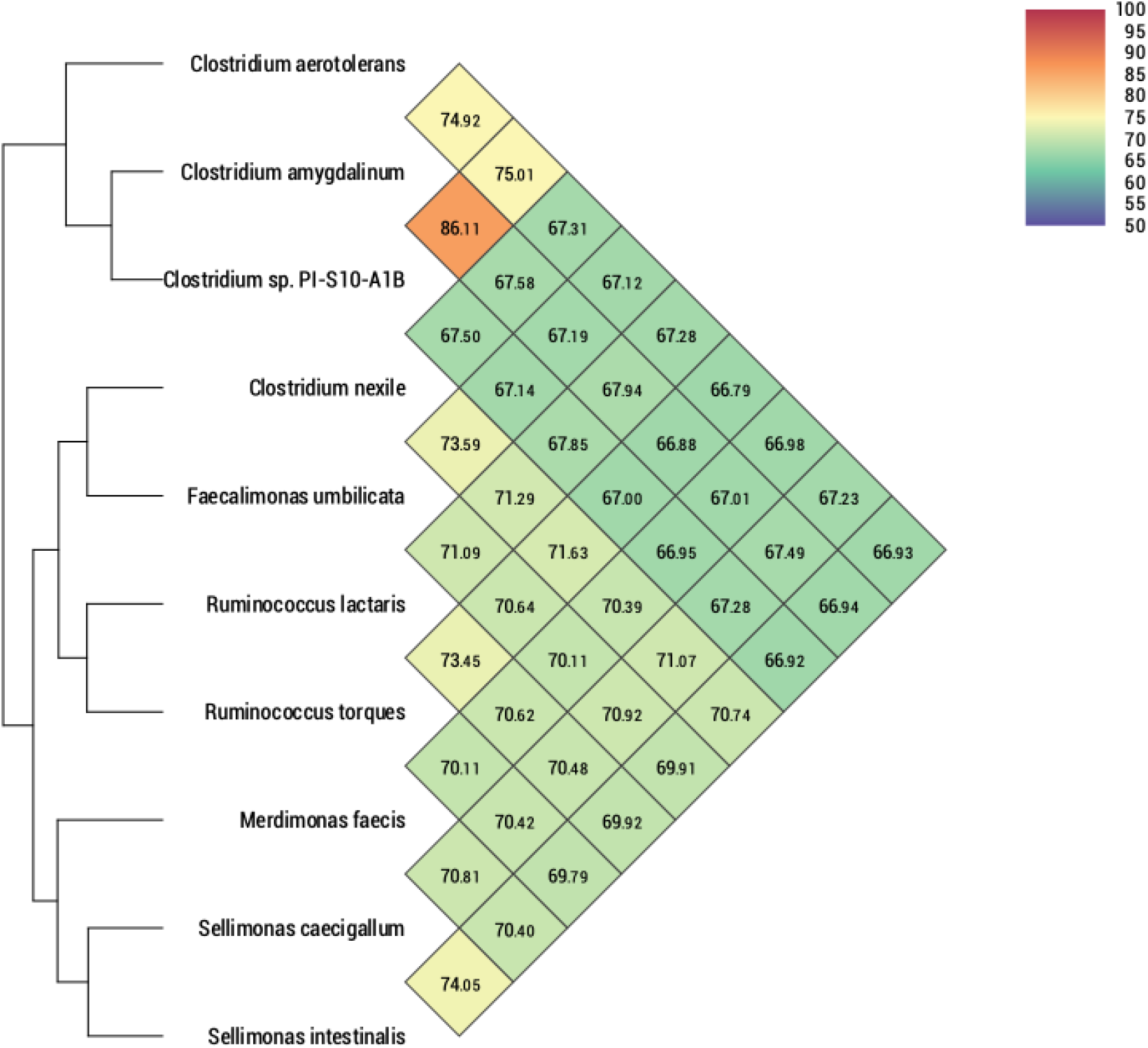
Average Nucleotide Identity comparison of Strain SW451 and related species with a valid taxonomy: Heatmap was generated with OrthoANI values calculated using the OAT software between *Sellimonas caecigallum* and other closely related species with standing in nomenclature.

## Phenotypic characteristics

Colony morphology of strain SW451 was determined after 2-3 days of incubation on BHI-M agar plates. Gram-staining was performed using a Gram-Straining kit set (Difco), according to the manufacturer’s instructions. Cell morphologies of cultures during exponential growth were examined by scanning electron microscopy (SEM). Aerotolerance was examined by incubating cultures for 2 days under aerobic and anaerobic conditions. Growth of strain SW451 at 4, 20, 30, 37, 40 and 55° C was determined. For pH range, the pH of the medium was adjusted to pH◻4–9 with sterile anaerobic solutions of 0.1◻M HCl and 0.1M NaOH. Motility of this microorganism was determined using motility medium with triphenyltetrazolium chloride (TTC) (21). The growth was indicated by the presence of red color, reduced form of TTC after it is absorbed into bacterial cell wall. Other biochemical tests including utilization of various substrates and enzyme activities were determined using the AN MicroPlate (Biolog) and API ZYM (bioMérieux) according to the manufacturer’s instructions. For cellular fatty acid analysis, strain SW451 was cultured in BHI-M medium at 37◻°C for 24◻h under anaerobic condition. Cellular fatty acids were obtained from cell biomass and analyzed by GC (Agilent 7890A) according to the manufacturer’s instruction of Microbial Identification System (MIDI) (22).

Cells of the strain SW451 were Gram-positive coccobacilli with the size of 1.0–1.5 μm (Fig. 3). Colonies on BHI-M agar were ivory yellow, raised and entire edge with 0.2-0.5 cm in diameter. Strain SW451 grew between 37° C and 45° C with optimum growth at 45°C. The optimum pH for the growth was 7, and growth was observed at pH◻6–7.5. The strain grew strictly under anaerobic condition, indicating that this strain is an obligate anaerobe. Strain SW451 utilized D-arabitol, D-fructose, L-fucose, D-galacturonic, palatinose, and rhamnose, and exhibited positive detection of alkaline phosphatase, leucine arylamidase, valine arylamidase, acid phosphatase, α-galactosidase, β-galactosidase, and α-glucosidase. Based on the results obtained in Biolog AN microplate and API ZYM, the carbon utilization and enzyme activity of strain SW451 were different from the closely related strains (Table 1). The predominant cellular fatty acids of strain SW451 included C _16◻:◻0_ (20.80%), C _14◻:◻0_ (17.98%) and C _16◻:◻0_ DMA (15.83%) in which differed from the reference strains (Table 2). The overall characteristics of the strain were summarized in Table 3.

**Table 1.** Characteristics of SW451 and closely related strains. Strains: 1, SW451; 2, *S. intestunalis* DSM 103502^T^; 3, *C. nexile* KCTC 5578^T^; 4, *M. faecis* KCTC 15482^T^. Results for metabolic end products of SW165 are from this study with cells that were cultured for 3 days at 37° C in BHI-M. +, positive; -, negative; w, weak; ±, variable; ND, not determined.

| Characteristic | 1 | 2 | 3 † | 4 ‡ |
| --- | --- | --- | --- | --- |
| Gram stain | + | + | + |  |
| Growth at 45 | + | + | + |  |
| Motility | - | - | ND | - |
| Carbon source (BIOLOG AN) |  |  |  |  |
| D-Arabitol | + | - | ND | ND |
| D-Fructose | + | + | ND | ND |
| L-Fucose | + | + | ND | ND |
| D-Galactose | - | + | ND | ND |
| D-Galacturonic | + | + | ND | ND |
| D-Glucose | - | + | + | - |
| D-Lactose | - | + | + | - |
| Maltose | - | + | - | - |
| Palatinose | + | + | ND | ND |
| Raffinose | - | + | - | ND |
| Rhamnose | + | + | ND | ND |
| Sucrose | - | - | + | ND |
| Enzyme activity (API ZYM) |  |  |  |  |
| Alkaline phosphatase | w | - | ND | + |
| Esterase (C4) | - | - | + | + |
| Leucine arylamidase | + | + | - | ND |
| Proline arylamidase | - | - | + | + |
| Valine arylamidase | w | + | - | ND |
| Alpha-chymotrypsin | - | w | - | ND |
| Acid phosphatase | w | + | ND | ND |
| Alpha-galactosidase | w | + | ND | + |
| Beta-galactosidase | + | + | ND | + |
| Alpha-glucosidase | + | + | ND | + |
| N-acetyl-beta-glucosaminidase | - | + | ND | ND |
| DNA G+C content (mol%) | 45.13 | 45.3 | 40.1 | 47 |

† Data from Seo et al. (2016)
‡ Data from Seo et al. (2017)

**Table 2.** Cellular fatty acid compositions of strains SW451 and related strains. Strains: 1, SW451; 2, *S. intestinalis* DSM 103502^T^; 3, *C. nexile* KCTC 5578^T^; 4, *M. faecis* KCTC 15482^T^. Values are percentages of total fatty acids detected. Fatty acids with contents of less than 1% in all strains are not shown; -, Not detected.

|  | 1 | 2 | 3 † | 4 ‡ |
| --- | --- | --- | --- | --- |
| **Straight chain** |  |  |  |  |
| C<sub>12:0</sub> | 3.62 | 2.45 | 5.6 | 3.4 |
| C<sub>14:0</sub> | 17.98 | 21.75 | 8.3 | 25 |
| C<sub>16:0</sub> | 20.80 | 27.45 | 36 | 18.6 |
| C<sub>16:0</sub> aldehyde | 7.03 | 1.94 | - | - |
| C<sub>18:0</sub> aldehyde | 1.50 | 0.6 | - | - |
| **Demethylacetal (DMA)** |  |  |  |  |
| C<sub>14:0</sub> DMA | 6.10 | 1.97 | - | - |
| C<sub>16:0</sub> DMA | 15.83 | 9.28 | - | - |
| C<sub>18:0</sub> DMA | 5.93 | 3.14 | - | - |
| **Unsaturated** |  |  |  |  |
| C<sub>13:1</sub> at 12-13 | 4.28 | 0.8 | - | 2.6 |
| C<sub>16:1</sub> *ω7c* | 2.35 | 4.41 | - | - |
| C<sub>18:1</sub> *ω7c* | 2.07 | 7.25 | - | - |
| C<sub>18:1</sub> *ω7c* DMA | 2.97 | 8.56 | - | - |
| Summed Feature 1 | 4.28 | 0.8 | 2.9 | 7.4 |
| Summed Feature 8 | 1.19 | 1.59 | 5.7 | 0.9 |
| Summed Feature 10 | 2.07 | 7.25 | - | - |

† Data from Seo et al. (2016)
‡ Data from Seo et al. (2017)
\*Summed features are fatty acids that could not be separated using the MIDI System. Summed feature 1 contains C<sub>13:1</sub> and/or C<sub>14:0</sub> aldehyde. Summed feature 13 contains C<sub>15:0</sub> anteiso and/or C<sub>14:0</sub> 2-OH; summed feature 8 comprises C<sub>17:1</sub> *ω8c* and/or C<sub>17:2</sub> at 16; summed feature 10 contains C<sub>18:1</sub> *ω7c* and/or unknown.

**Table 3.** Description of strain SW451, according to the digitized protologue (www.imedea.uib.es/dprotologue) website.

|  |  |
| --- | --- |
| Taxonumber | TA01013 |
| Species name | Sellimonas caecigallum |
| Genus name | Sellimonas |
| Species epithet | caecigallum |
| Species status | sp. nov. |
| Species etymology | ce.ci.gal'lum N.L. gen. n. referring to the caecum of a chicken where the type strain was isolated. |
| Designation of the type strain | SW451 |
| Strain collection numbers | DSM 109473 |
| 16s Rrna gene accession number | MN053974 |
| Genome accession number | PRJNA551641 |
| Genome size | 2.67 Mbp |
| GC mol% | 45.13 |
| Country of origin | United State |
| Region of origin | Brookings, South Dakota |
| Source of isolation | caecum of feral chickens |
| Gram stain | positive |
| Cell shape | small chain coccobacilli |
| Cell size | 1.0–1.5 µm |
| Motility | non-motile |
| Colony morphology | 0.2–0.5 cm in diameter, ivory yellow, raised with entire edge |
| Temperature range | 37–45° C |
| Temperature optimum | 45 °C |
| pH range for Growth | 6–7.5 |
| pH optimum | 7 |
| Relationship to O<sub>2</sub> | anaerobe |

**Fig. 3.**
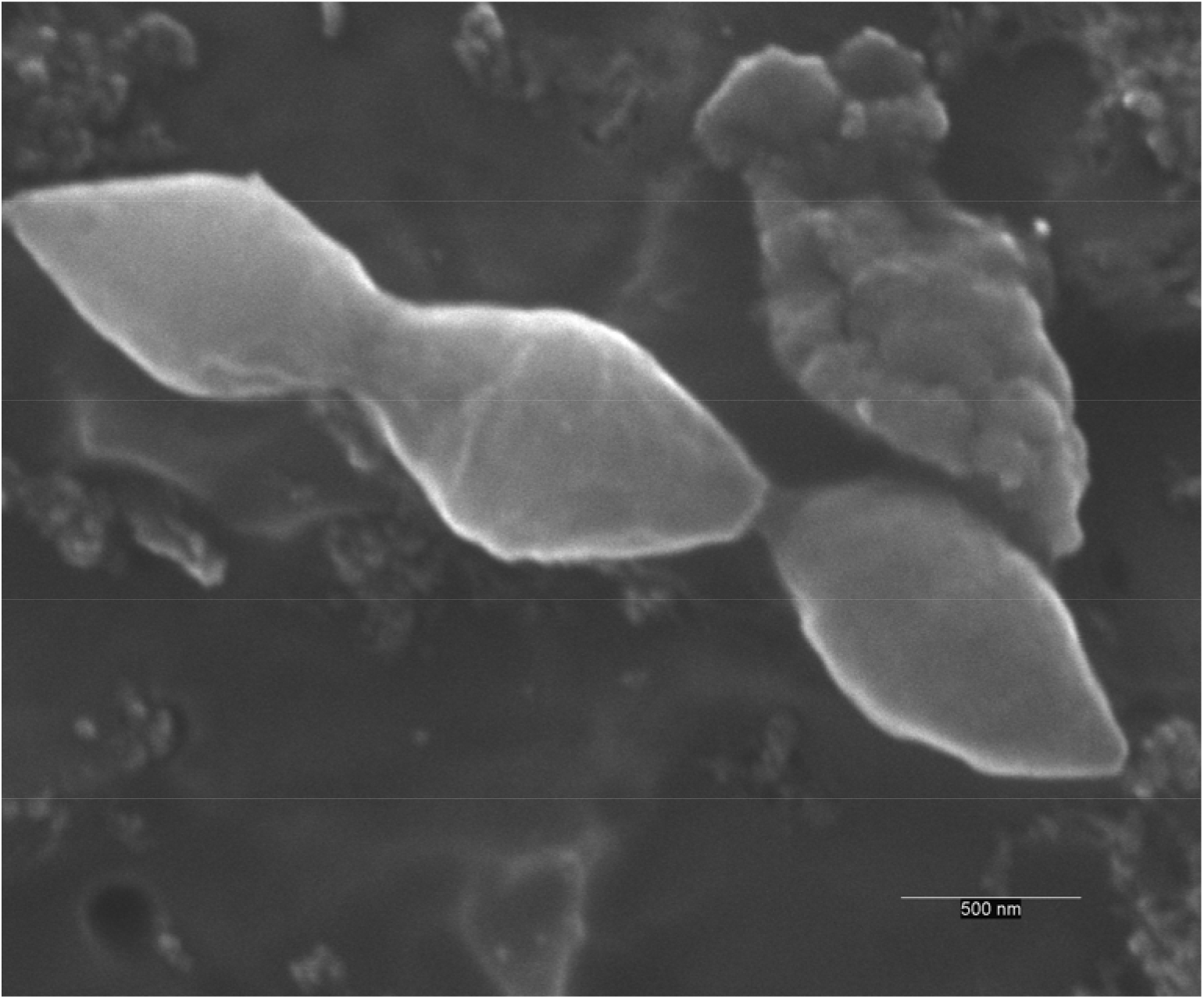
Scanning electron micrograph of cells of SW451. Cells were anaerobically cultured for 1 day at 37° C in BHI-M medium. Bar, 2 μm.

## Description of *Sellimonas caecigallum* SW451 sp. nov

*Sellimonas caecigallum* sp. nov. (referring to L. n. *caecum*, caecum; L. gen. n. *gallum*, of a chicken; N.L. gen. n. *caecigallus*, from the caecum of a chicken). Cells are strictly anaerobic, Gram-strain-positive and non-motile. The average size of each cell is 1.0–1.5 μm and coccobacillus-shaped. Colonies are visible on BHI-M agar after 2 days and are approximately 0.2–0.5 cm in diameter, ivory yellow, raised with entire edge. The strain exhibits optimal growth in BHI-M medium at 45° C and pH 7. The strain utilizes D-arabitol, D-fructose, L-fucose, D-galacturonic, palatinose, and rhamnose as a carbon source. Positive enzymatic reactions are obtained for alkaline phosphatase, leucine arylamidase, valine arylamidase, acid phosphatase, α-galactosidase, β-galactosidase, and α-glucosidase. The primary cellular fatty acids are C _16◻:◻0_, C _14◻:◻0_ and C _16◻:◻0_ DMA. The genome of this strain is 2.67 Mbp with 45.13 mol% of G+C content. isolated from the cecum of feral chicken. The type strain is SW451 (DSM 109473^T^)

## Nucleotide sequence accession number

The 16S rRNA gene and genome sequence were deposited in GenBank and SRA under accession numbers MN053974 and PRJNA551641, respectively.

## Deposit in culture collection

Strain SW451 was deposited in the DSMZ culture collection under number DSM109473.

## Acknowledgements

This work was supported in part by the USDA National Institute of Food and Agriculture, Hatch projects SD00H532-14 and SD00R540-15, and a grant from the South Dakota Governor’s Office of Economic Development awarded to JS. SW received support from the Science Achievement Scholarship of Thailand.

